# Domain Dissolution in Supported Lipid Bilayers Triggered by Unsaturated Phospholipid Addition

**DOI:** 10.64898/2026.05.20.726269

**Authors:** Adeyemi T. Odudimu, Nathan J. Wittenberg

## Abstract

Significant cellular processes, including protein sorting, signal transduction, and pathogen entry, amongst others, are associated with membrane microdomains, also known as lipid rafts. Lipid rafts, due to their unique biophysical properties compared to their surrounding environment, which stem from their distinct lipid and protein profiles, have garnered interest in methods and techniques that tune their coexisting liquid-ordered/liquid-disordered state, aiming to disrupt or destabilize them. Since cholesterol stabilizes the membrane domain, cholesterol-depleting compounds like cyclodextrin can be used to destabilize and disrupt the membrane rafts. Overall, given the membrane raft’s importance in biological processes, it is crucial to understand the biophysical factors that influence its stability. In this study, we present a new method for disrupting and dissolving lipid rafts in a model system of phase-separated supported lipid bilayer (SLB) patches composed of DOPC, DPPC, and cholesterol. Using fluorescence microscopy to monitor the liquid ordered (Lo) and liquid disordered (Ld) phases of the SLB patches, we observed that adding DOPC liposomes causes a transformation of the co-existing Ld and Lo phases into a single-phase bilayer. On the other hand, adding liposomes that match the lipid content of the phase-separated SLB patch increase the areas of the existing Ld and Lo phases. This work also offers a new method for redistributing raft-localized molecules, confirmed by tracking the redistribution of cholera toxin bound to GM1 after domain dissolution with DOPC liposomes. The work describes an alternative method for dynamically altering membrane composition and dissolving domains via liposome addition, rather than lipid depletion or exchange.

## INTRODUCTION

Lipid rafts, also known as membrane microdomains or nanodomains, depending on their size, are heterogeneous zones of the cellular plasma membrane.^1^ Lipid rafts, originally recognized due to their resistance to detergents, are enriched in saturated phospholipids, sphingolipids, and cholesterol, which form an ordered subcompartment on the plasma membrane. Compared to their more disordered surroundings, which are enriched in unsaturated lipids, lipid rafts can serve as organizing centers for membrane proteins that enable their specialized cellular functions.^2,3^

On cell surfaces, lipid rafts play a significant role in signal transduction pathways by serving as reservoirs for signaling proteins and receptors.^4,5,6^ Lipid rafts are also associated with immune signaling.For example, T-cell weakening viruses, such as HIV, use lipid rafts as a point of entry into host cells via its CD4 receptor localization.^7,8^ Due to their association with membrane trafficking processes, such as endocytosis, lipid rafts are involved in the transport of molecules across the plasma membrane.^9^ Moreover, there is significant evidence linking various diseases with lipid raft-associated proteins.^10^ Furthermore, protein receptors that co-localize with lipid rafts undergo significant alterations in various physiological conditions, including viral and microbial infections, cardiovascular disorders, neurological diseases, and cancer.^11,12,13^

Additionally, the cellular immune response against pathogens and toxins is activated by signaling proteins and receptors localized in lipid rafts.^14,15,16,17^ Hence, pathogens have developed a potent strategy to evade immune responses during their pathogenesis by hijacking signaling proteins within membrane rafts,^18,19^ for example, coronavirus (SARS-CoV-2) receptor (ACE-2) had been identified to bind to glycosphingolipid in the host’s lipid raft in the process of its entry into the cell,^20,21^ this is not only peculiar to SARS-CoV-2 but also to other viruses and pathogens.^22,23,24,25^ This has led to a strategy for developing therapeutics against these pathogens and other lipid-raft-associated diseases.^26,27^ Disruption of lipid rafts by cholesterol-inhibiting drugs or cholesterol-extracting agents, including methyl-β-cyclodextrin (MβCD),^28,23^ statins, ^29^ filipin,^30^ saponin,^31^ lovastatin,^32^ and squalestatin^33^, has been reported to have antiviral activity.

Because lipid rafts and raft-associated biomolecules are crucial for essential biological processes, the lipid composition that supports the formation of lipid rafts is tightly regulated. The key constituents of the membrane that support raft formation, such as unsaturated and saturated phospholipids, sphingolipids, and cholesterol, have relative mole fractions that fall within ranges that support spontaneous phase separation under certain conditions.^34^ To probe the implications of alterations to the relative proportions of membrane lipids, researchers have turned to techniques involving selective lipid extraction or lipid exchange,^35^ typically enabled by various cyclodextrins. A common technique involves selective cholesterol extraction or delivery by MβCD,^28,36^ however, using specialized cyclodextrins, phosphatidylcholines and sphingomyelin can be exchanged. Being an exchange process, the delivery of an exogenous lipid is accompanied by the extraction of an endogenous lipid.

The consequences of lipid extraction or exchange have been investigated in a variety of model systems, including giant unilamellar vesicles (GUVs), large unilamellar vesicles (LUVs), liposomes, and supported lipid bilayers (SLBs).^37,38,39,40^ Model lipid membrane systems are advantageous because they can be prepared with precisely-defined compositions, offering a level of compositional control not available in cellular systems.^41^ In these systems, mixing saturated and unsaturated phospholipids with cholesterol in the right proportions can result in the membrane separating into distinct liquid ordered (Lo) and liquid disordered (Ld) phases, where the Lo phase mimics lipid rafts and is enriched in saturated phospholipids and cholesterol.^42,43^ Over the years, a number of groups have investigated the formation and properties of Lo and Ld phases, and the use of these model systems enables the examination of how dynamic changes in lipid composition alters Lo and Ld domains. For example, Veatch and Keller, using GUVs, reported that a lipid mixture of saturated and unsaturated lipids, and cholesterol at varying concentrations and temperatures resulted in a coexisting Lo and Ld domains, which can be tuned by temperature changes and cholesterol depletion using β-cyclodextrin.^44^ Also, Carrer and co-workers in their work synthesized cholesterol derivatives that can disrupt the membrane phase of GUVs via a detergent-like effect.^45^ Lipid extraction via selective binding of amphiphilic Janus nanoparticles to Ld phase of GUVs has also been shown to result in microdomain destabilization and shrinkage.^46^

In contrast to previous studies, here we describe a method to deliver liposomes with defined compositions to SLB patches as a means to dynamically alter membrane composition. Our method bears some resemblance to the approach developed by Enoki and Feigenson, where GUV lipids are exchanged with SLB lipids,^47^ but we expect that all liposomal lipids to be fully incorporated into the SLB. Importantly, our method is neither a lipid extraction nor exchange method, but rather it is strictly a lipid addition method that alters the mole fraction of lipids globally within the SLB patch. In doing so, the lipid composition of the SLB patch shifts toward the composition of the added liposomes. This is a fundamentally different compositional shift than is possible with lipid extraction or exchange methods. We show that delivery of liposomes rich in unsaturated phospholipids to SLB patches with preexisting Lo and Ld domains triggers a miscibility transition that is caused by enrichment in the patch in unsaturated phospholipids and is accompanied by a relative decline in the mole fraction of saturated phospholipids and cholesterol. Conversely, delivery of liposomes with compositions that match that of the phase-separated SLB patch preserves and enlarges coexisting Lo and Ld domains. Finally, we show that addition of unsaturated lipids to phase-separated SLB patches causes the redistribution of cholera toxin bound to the Lo-associated ganglioside GM1.

## MATERIALS AND METHODS

### Reagents and Chemicals

Dioleoylphosphatidylcholine (DOPC), dipalmitoylphosphatidylcholine (DPPC), Cholesterol (Chol), and 23-(dipyrrometheneboron difluoride) - 24 - norcholesterol (TF-Chol) were obtained from Avanti Polar Lipids (Alabaster, AL). Texas Red - 1,2 - dihexadecanoyl - sn - glycero - 3 - phosphoethanolamine, trimethylammonium salt (TR-DHPE) and tris(hydroxymethyl)aminomethane (Tris base) were purchased from Fisher Scientific. Sucrose, sodium chloride, calcium chloride, and chloroform were purchased from Sigma Aldrich.

### Preparation of GUVs

Both single and phase-separated GUVs were prepared using the electroformation swelling method.^48^ For the phase-separated GUVs, lipid mixtures of DOPC, DPPC, and cholesterol in chloroform with ratios corresponding to the compositions along the tie line^49^ **(Figure 2a)** were prepared to a total concentration of 1 mg/mL and vacuum-dried for at least 2 hours to form a lipid film. To mark the liquid-ordered (Lo) and liquid-disordered (Ld) phase, 0.5 and 1 mol% of TR-DHPE and TF-Chol were added, respectively. When it was included, GM1 was 1 mol %. Single-phase GUVs were prepared using 99.5% DOPC and 0.5% TR-DHPE. The film was resuspended in 60 μL of chloroform and vortexed. Then, 22 μL of the solution was continuously transferred to form a thin film on an indium tin oxide (ITO) slide that had been cleaned with 2% Alconox and water. Chloroform was evaporated in a vacuum desiccator. Using a second ITO slide, a capacitor was formed, and 350 μL of 200 mM sucrose was added as a swelling agent. A waveform generator was used to supply a 10 Hz AC signal with 10 volts peak-to-peak across the ITO slides. The capacitor was placed in an oven for at least 2 hours at 60 °C. The electroformed GUVs were removed with a pipette and kept in an amber Eppendorf tube until further use.

### Formation of SLB Patches

To convert the spherical GUV into a planar SLB patches, first, round coverslips were cleaned with a flow of nitrogen gas to remove dust. This was followed by rinsing with isopropyl alcohol and Milli-Q water. After drying, the coverslips were soaked in 2% sodium dodecylsulfate (SDS) for at least 30 minutes, thoroughly rinsed with Milli-Q water, then blown dry with nitrogen gas. Finally, the coverslips were cleaned in a UV-Ozone chamber (ProCleaner Plus, Bioforce Nanosciences). GUVs were diluted to a total lipid concentration of 3 μg/mL in Tris buffer with calcium (10 mM Tris base, 150 mM NaCl, and 2.2 mM CaCl_2_, pH 7.0). The GUV solution was introduced onto the precleaned microscope slide mounted in an AttoFluor cell chamber. Using an incubation time of 2 minutes, the GUVs ruptured on glass to form SLB patches. Unruptured GUVs were washed away with calcium-free Tris buffer (10 nM Tris base and 150 nM NaCl, pH = 7.0).

### Preparation of liposomes

Chloroform solutions containing 1 mg/ml DOPC were dried under vacuum for at least 2 hours to form a lipid film. The film was resuspended in Tris buffer with calcium and vortexed until thoroughly mixed. The hydrated solution was bath-sonicated for 10 minutes at room temperature using a Branson 3510 ultrasonic cleaner. Finally, the solution was extruded by passing through a 50 nm polycarbonate membrane filter 23 times in a Mini-Extruder (Avanti). The stock liposome solution was diluted to concentrations of 0.05, 0.1, and 0.2 mg/ml depending on the experiment. In some experiments, liposomes were prepared from 0.1 mg/ml of 2:2:1 DOPC/DPPC/Chol, and 4:1 DOPC/Chol. All preparation steps for these vesicles were done at 45 °C.

### Fluorescence Imaging of SLB Patches

Fluorescence imaging was performed using an inverted microscope (Eclipse Ti, Nikon), controlled by Nikon Elements software, and equipped with a 100× oil immersion objective featuring a 1.49 numerical aperture. The appropriate filter sets were used to capture the Ld phase, labeled with TR-DHPE, and the Lo phase, labeled with TF-Chol. All images and movies were acquired using fluorescence excitation with an LED light engine (Aura II, Lumencor) and a sCMOS camera (Orca Flash 4.0 v2, Hamamatsu) at a 512 × 512 pixel field of view. Images were processed using ImageJ Fiji software.

### Fluorescence Recovery After Photobleaching (FRAP)

The diffusion coefficient of lipids in the SLB patches pre- and post-liposome addition was determined by FRAP analysis using an inverted microscope. Using a 2 s 405 nm (50 mW) laser pulse, a region of interest (ROI) was photobleached on the Ld phase labeled with TR-DHPE, and fluorescence recovery was captured for 60 s at 1 s intervals. The images were analyzed to determine the diffusion coefficient using the Hankel transform method described by Jonsson and coworkers.^50^

### Data Analysis

Graphs and data analysis were performed using GraphPad Prism 8 software. All experiments were done with a minimum of three replicates.

## RESULTS AND DISCUSSION

### Formation of SLB patches

To form planar SLB patches, first, GUVs were prepared by electroformation using ternary mixtures of DOPC, DPPC, and cholesterol, the molecular structures of which are shown in **Figure 1**. The specific mole fractions of DOPC, DPPC, and cholesterol used to prepare the GUVs are shown in the phase diagram in **Figure 2a**. These compositions included 100 % DOPC (composition *i*), as well as mixtures with DOPC/DPPC/cholesterol mole fractions of 54:30:16 (composition *ii*), 2:2:1 (composition *iii*), and 19:55:25 (composition *iv*). Compositions *ii, iii*, and *iv* are all within the 2-phase (Lo and Ld) coexistence region of the phase diagram,^51^ whereas composition *i* is solely DOPC and thus completely Ld. Additionally, compositions *i, ii*, and *iii* lie along a tie line,^49^ therefore the Lo and Ld phases of these mixtures have the same compositions. However, compositions *i, ii*, and *iii* differ in their relative area fractions of the Lo and Ld phases.

**Figure 1:**
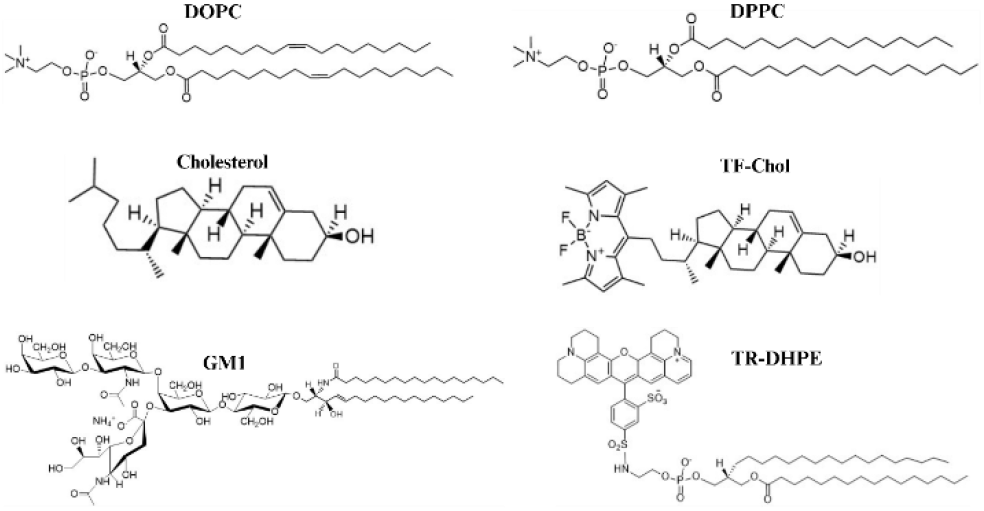
Molecular structures of the lipid molecules used in this study.

**Figure 2:**
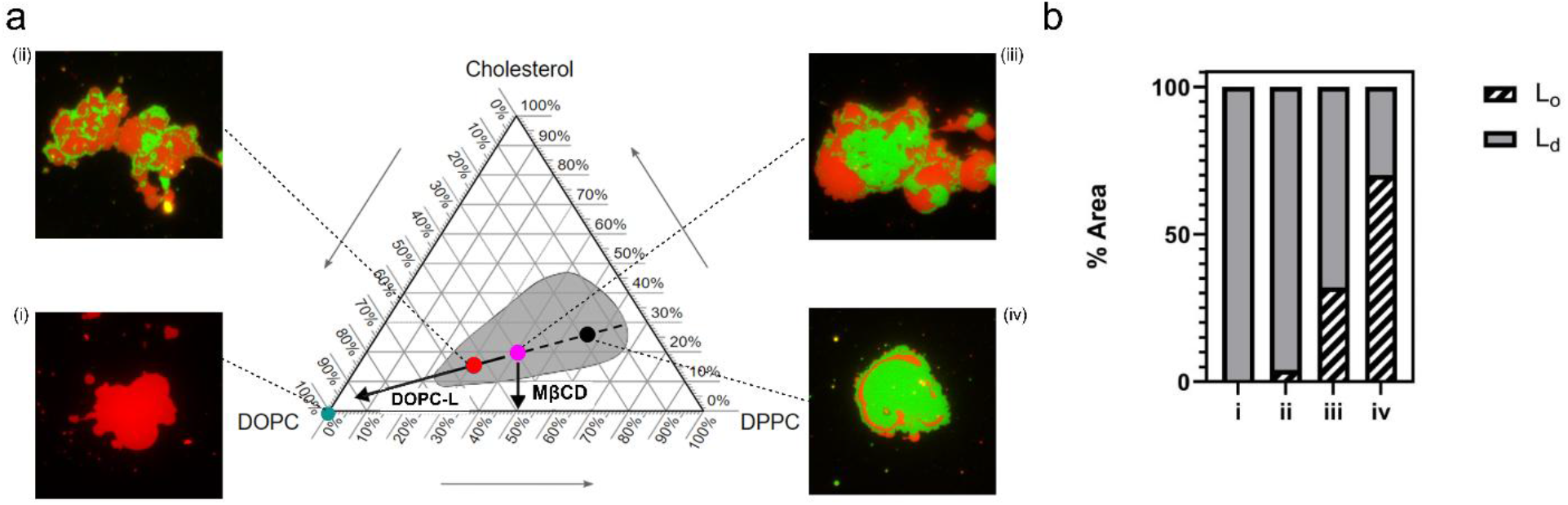
SLB patch lipid compositions and Lo and Ld area fractions. (a) Phase diagram of ternary mixtures of DOPC/DPPC/Chol, where compositions with coexisting Ld and Lo phases are shaded in gray. The teal, red, magenta, and black dots represent compositions *i, ii, iii*, and *iv* shown in the fluorescence images. The fluorescence images ae SLB patches where the red areas are Ld domains labeled with TR-DHPE, and the green areas are Lo domains labeled with TF-Chol. The diagonal (DOPC-L) and vertical (MβCD) arrows depict how membrane composition changes with DOPC addition by liposome fusion at SLB patch edges and cholesterol extraction by MβCD, respectively. (b) Area fractions of Ld and Lo phases in SLB patches with compositions *i, ii, iii*, and *iv*.

To evaluate the area fractions of Lo and Ld phases in the three phase-separating mixtures, after GUVs were formed, they were allowed to settle to the bottom of the sample chamber where they promptly ruptured onto the glass substrate to form a SLB patch. To facilitate GUV rupture calcium (2.2 mM) was included in the buffer. In these GUVs, the Ld phase was marked with Texas Red-DHPE (TR-DHPE) and the Lo phase was marked with TopFluor-cholesterol (TF-Chol). As expected, composition *i* did not exhibit phase separation. Also, as expected, the area fraction of the Lo domains in compositions *ii, iii*, and *iv* increased as the mole fractions of DPPC and cholesterol increased. Example images of SLB patches with these compositions are shown in **Figure 2a**, while measured average area fractions of the Lo and Ld domains are shown in **Figure 2b**. The diagonal arrow labeled DOPC-L in **Figure 2a** depicts the compositional shift associated with addition of DOPC liposomes to SLB patches. In contrast, the downward vertical arrow represents the how the lipid composition of a SLB patch with composition *iii* would be altered upon cholesterol extraction by MβCD. In both cases, the compositional shifts push the overall compositions out of the two-phase coexistence region (gray). However, the means by which lipid miscibility is triggered (DOPC addition vs. cholesterol depletion) differ significantly.

### Addition of liposomal lipids to SLB patches

To investigate how liposomal lipids add to a SLB patch, we started with a single-phase SLB patch composed of DOPC, composition *i* in **Figure 1a**, labeled with TR-DHPE and introduced unlabeled DOPC liposomes. **Figure 3** illustrates the experiment. SLB edges are energetically unstable^52^ and are favorable sites for liposome adsorption,^53^ which adds membrane material to the SLB patch and reduces the ratio of membrane edge length to surface area. Reducing this ratio shifts the balance of energetically unfavorable membrane edges toward more favorable SLB-substrate adhesive interactions.^54^ Because the DOPC liposomes are unlabeled, their adsorption and rupture on the substrate is invisible, when in isolation. However, when the unlabeled liposomes rupture at the edge of the SLB patch they add to the patch area. Because the new DOPC from the liposomes becomes part of the SLB patch, the TR-DHPE from the patch spreads into the added membrane area.

**Figure 3:**
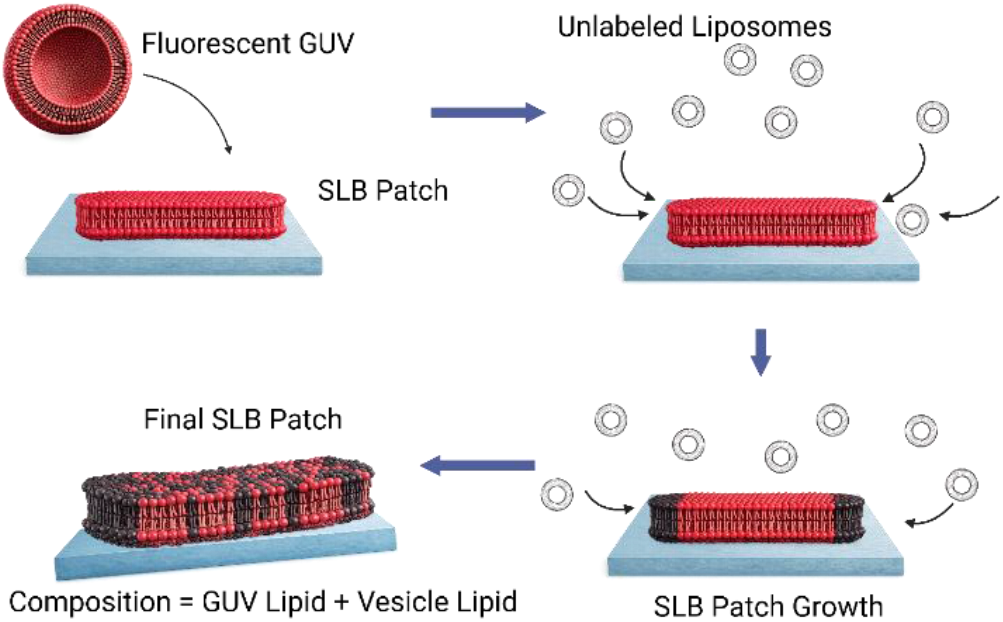
Illustration of SLB expansion on the addition of DOPC liposomes. First, a single-phase GUV (red sphere) adsorbs and bursts on the glass substrate to form an SLB patch. Then, DOPC liposomes (black bilayer circles) adhere to the substrate and rupture at the edge of the phase-separated SLB patch. As the process proceeds, the DOPC from the liposomes becomes incorporated into the SLB patch, which enriches the patch with more DOPC, resulting in a larger SLB.

To observe SLB patch spreading in response to the addition of unlabeled DOPC liposomes, we conducted time-lapse fluorescence microscopy. **Figure 4** shows results from these experiments. Prior to the addition of DOPC liposomes (t = 0 s), the SLB patch exhibits rather smooth curved edges and uniform fluorescence intensity over its area. If no DOPC vesicles are added, the SLB patch is stable and does not spread. A 100 µL volume of 0.1 mg/mL DOPC liposomes were added to the chamber with the SLB patches. Shortly after addition of the DOPC liposomes (t = 100 s), the SLB patch begins to enlarge. This is due to TR-DHPE spreading into the newly formed SLB membrane along the patch edge. Additionally, satellite patches adjacent to the largest patch begin to enlarge. Patch enlargement is a continuous process, and as it continues (t = 180 s), the largest SLB patch eventually subsumes the satellite patches. Notably, as the SLB patch grows, the edge becomes more jagged. A closer examination of the edge growth process reveals this jagged geometry **(Figure S1)**. This occurs because liposome rupture and membrane addition at the patch edge occurs in a spatially-random manner. In some respects, this process resembles the spontaneous spreading of intentionally delaminated SLBs,^55^ however, the process does not cease because new lipid material is continuously added to the SLB edge. As the SLB patch grows, the edges appear dimmer than the center because the distance for TR-DHPE to diffuse into the newly formed membrane is continuously increasing. Approximately five minutes after DOPC liposome addition (t = 300 s), the SLB patch has filled the field of view, but the remnants of the original patch are faintly observable as an area of slightly greater fluorescence intensity. Eventually, the TR-DHPE fluorescence spreads uniformly over the entire field of view (t = 500 s).

**Figure 4.**
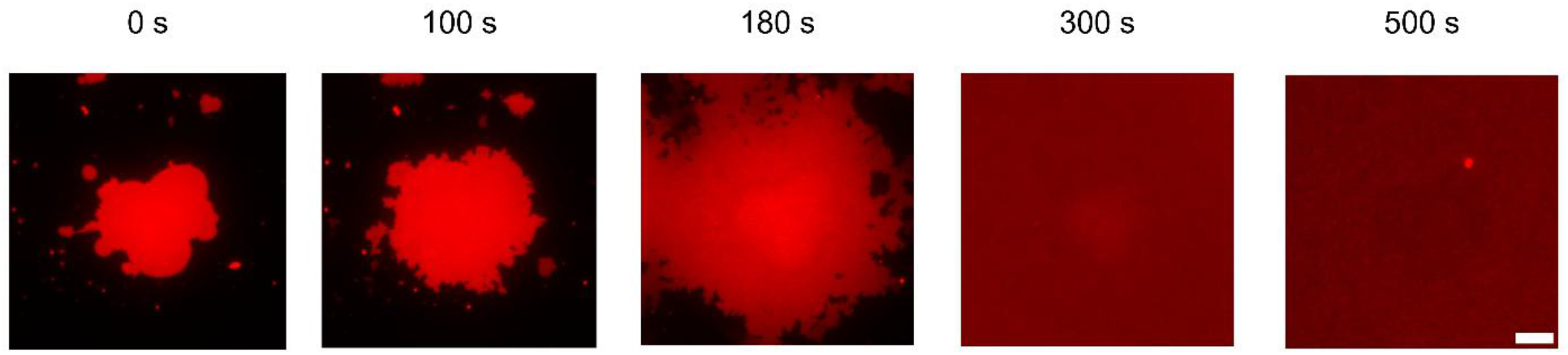
Fluorescence micrographs showing the effect of the addition of 100% DOPC liposomes to a single-phase SLB patch with a composition of 99.5:0.5 DOPC/TR-DHPE. Scale bar: 5 µm.

### Shifting SLB patch composition

The experiment shown in **Figure 4**, where the SLB patch and liposomes have the same lipid composition, demonstrates that liposomal lipids can be readily incorporated into a SLB patch. These observations set the stage for experiments where the SLB patch and liposomes have different compositions, and the incorporation of liposomal lipids dynamically shifts the composition of the SLB patch. In these experiments, we started with phase-separated SLB patches prepared from GUVs as described above. The first composition we tested had mole fractions of DOPC/DPPC/Chol of 2:2:1, and the Ld and Lo phases were labeled with TR-DHPE and TF-Chol, respectively. This lipid ratio corresponds to composition *iii* in **Figure 1**. Upon formation of the SLB patch, unlabeled DOPC vesicles were introduced, as shown in **Figure 5**.

**Figure 5:**
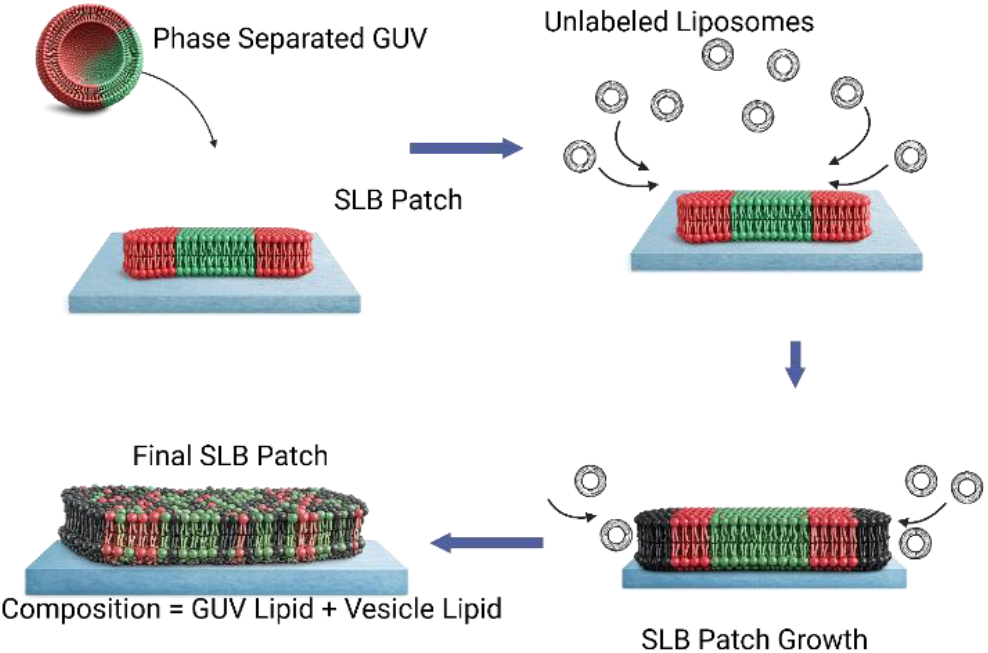
Illustration of domain dissolution by DOPC liposomes. First, a phase-separated GUV (green and red sphere) adsorbs and bursts on the glass substrate to form a SLB patch. Then, DOPC liposomes (black bilayer circles) adhere to the substrate and rupture at the edge of the phase-separated SLB patch. As the process proceeds the DOPC from the liposomes becomes incorporated into the SLB patch, which enriches the patch in DOPC. Eventually, the composition of SLB patch becomes highly enriched in DOPC, triggering miscibility of the DOPC, DPPC, and cholesterol and the disappearance of distinct Lo and Ld domains.

As the DOPC vesicles add to the SLB patch edge the overall fraction of DOPC in the patch increases, and at the same time the fractions of DPPC and cholesterol decrease. This should result in the composition of the patch shifting to outside of the two-phase coexistence region and dissolution of the Lo and Ld domains, as illustrated by the diagonal arrow in **Figure 2a. Figure 6** shows still images, and **Movie S1** shows a time-lapse video of the entire process. Prior to addition of DOPC vesicles, the Lo and Ld domains are distinctly separated. Then, a 100 µL aliquot of 0.1 mg/mL DOPC liposomes were added to the chamber with the SLB patches. After one minute (t = 60 s) of DOPC vesicle exposure, the SLB clearly enlarges and both the TR-DHPE and TF-Chol markers spread along the patch periphery. At this time, however, the size and shape of the Lo domains are relatively unchanged. As time elapses and more DOPC is added to the SLB patch edge, the size and shapes of the domains clearly change. At t = 130 s, the Lo domains are shrinking, and the smaller satellite Lo domains disappear entirely. At the same time the spread of both the Lo and Ld markers (TF-Chol and TR-DHPE) into the newly formed SLB are apparent. At an even longer time point (t = 220 s), the beginnings of greater lipid miscibility can be seen. The Lo areas have significantly shrunk, with a faint area of higher TF-Chol intensity in the remnant of the initial Lo domain, accompanied by TF-Chol and TR-DHPE spreading throughout the growing patch. At even longer time points (t = 1200 s), the TF-Chol and TR-DHPE fluorescence is uniform over the entire field of view, suggesting complete dissolution of the Lo and Ld domains. This transition to total miscibility is due to the increasing fraction of DOPC in the SLB patch, which pushes the lipid composition out of the two-phase coexistence region of the phase diagram.

**Figure 6:**
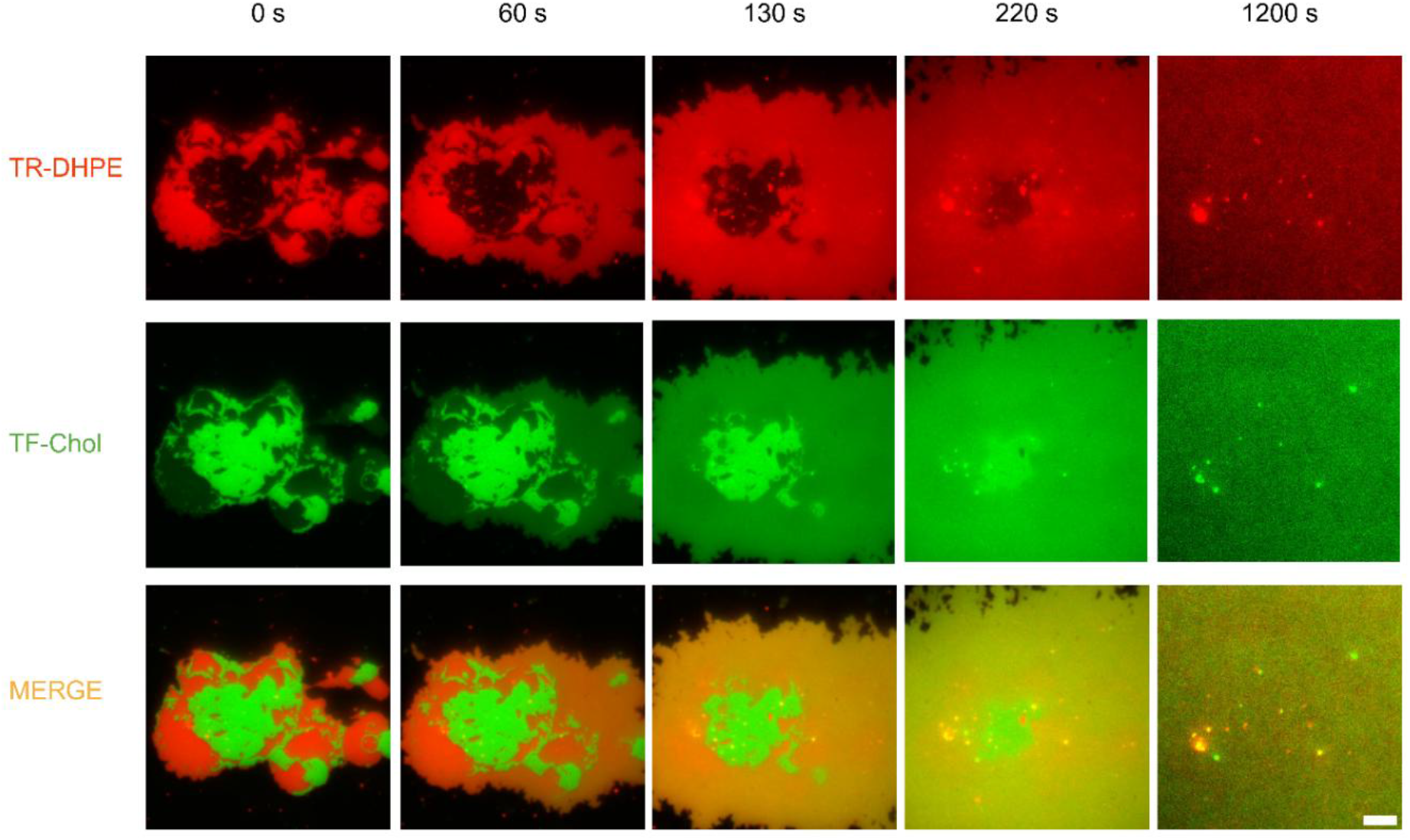
DOPC liposome interactions with 2:2:1 DOPC/DPPC/Chol phase-separated SLB patch. Fluorescence images showing how the addition of unlabeled DOPC liposomes the change of the morphology of phase-separated SLB patch (2:2:1 DOPC/DPPC/Chol) to form a single phase SLB. The top, middle, and bottom images represent the Ld phase labeled with TR-DHPE (red), Lo phase labeled with TF-Chol (green), and the merger of the two channels. The corresponding movie is in **Movie S1** (Supporting Information). Scale bar: 5 µm.

We also explored whether the relative area fractions of the Lo and Ld domains in SLB patches had an influence on whether or not the addition of DOPC liposomes caused domain dissolution. To reduce the area fraction of the Lo domains and enlarge the Ld domains, we used a lipid composition that was more rich in DOPC with reduced fractions of DPPC and cholesterol (DOPC/DPPC/Chol 54:30:16), which corresponds to composition *ii* in **Figure 1**. These SLB patches had a Lo area fraction of 4% **(Figure 1b)**. On the other hand, increasing the mole fractions of DPPC and cholesterol resulted in a gain in area fraction of the Lo domains in and a decrease in the area fraction of the Ld domains in SLB patches. This composition had a DOPC/DPPC/Chol molar ratio of 19:55:25 and corresponds to composition *iv* in **Figure 1a**, which had an Lo area fraction of 68% **(Figure 1b)**. In both cases, the addition of DOPC liposomes caused complete dissolution of the distinct Lo and Ld domains **(Figures S2 and S3** and **Movies S2 and S3)**. This demonstrates that regardless of starting composition of the SLB patch, the lipids from DOPC liposomes readily add to the SLB patches, and enough DOPC can be added to push the compositions out of the 2-phase coexistence region of the phase diagram. This is especially noteworthy for the DOPC/DPPC/Chol 19:55:25 composition because significantly more DOPC must be added to shift the composition into the region of lipid miscibility.

### Effect of DOPC liposome concentration on domain dissolution

Since addition of DOPC liposomes caused dissolution of the distinct membrane domains in SLB patches, we reasoned that the rate at which the domains disappear would be related to the rate of DOPC liposome delivery, and thus the rate at which the DOPC mole fraction rises. A number of reports have shown that the concentration of liposomes has significant effect on the rate of SLB formation, with higher concentrations leading to accelerated SLB formation.^56^ Therefore, we predicted that changing the DOPC liposome concentration should alter the rate of domain disappearance. To examine this effect, we varied the DOPC liposome concentration added to the SLB patch chamber by injecting 100 µL aliquots with DOPC concentrations ranging from 0.05 mg/mL to 0.2 mg/mL.

To determine the rate of domain disappearance upon DOPC liposome addition, we monitored the fluorescence intensity of TF-Chol over time in Lo domains in SLB patches with a 2:2:1 DOPC/DPPC/Chol composition. TR-DHPE was also included in the SLB patches to locate the Ld domains and patch edges. Because the fluorescence intensity of TF-Chol varies somewhat from patch to patch, we normalized the intensity using the following formula:

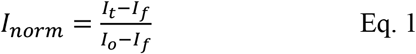

Where *I*_*o*_ is the initial TF-Chol fluorescence intensity in a ROI encompassing a Lo domain before DOPC liposome addition, *I*_*t*_ is the TF-Chol fluorescence intensity in the same ROI at time *t* after DOPC liposome addition, and *I*_*f*_ is the final TF-Chol fluorescence in the ROI after complete Lo domain disappearance. Like the results shown in **Figure 6**, once DOPC liposomes began adding to the SLB patch, the patch area increased and the distinct Lo and Ld domains disappeared. During this time, the normalized intensity of TF-Chol in the Lo domains decreased. When plotted relative to time, it is clear that the increasing the DOPC liposome concentration causes a sharp increase in the rate of TF-Chol intensity decline **(Figure 7)**. This confirms that the rate of domain disappearance is tightly linked to the rate at which lipids are added to the SLB patch perimeter, and thus the rate at which the global patch composition changes. It should also be noted that the lag time between DOPC liposome addition and pronounced TF-Chol intensity decline is inversely correlated with concentration. This agrees with many studies that show that SLB formation by liposome rupture is preceded by intact liposome accumulation on the substrate until a critical surface coverage is reached.^57,58,59^ The time to achieve critical surface coverage has been shown to be inversely correlated with liposome concentration,^60^ in agreement with our observations.^56^

**Figure 7:**
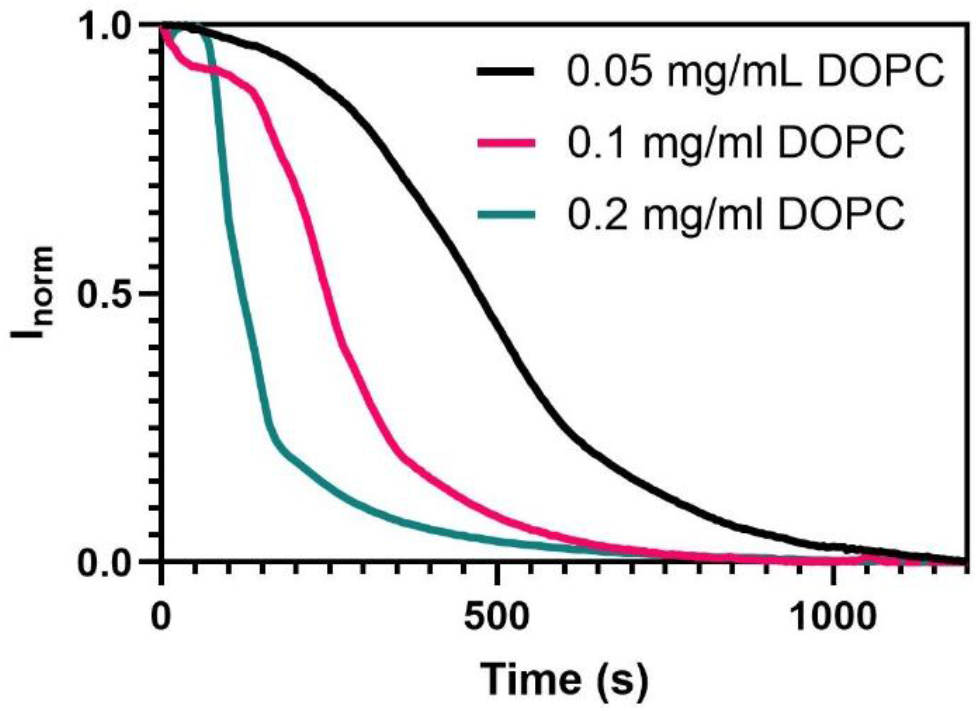
Effect of DOPC liposome concentration on rate of domain dissolution. The fluorescence intensity of TF-Chol in Lo domains declines as the domain disappears upon addition of DOPC liposomes. As the DOPC liposome concentration increases, the rate of TF-Chol fluorescence decline increases.

### Domain Disappearance is Accompanied by Changes in Lipid Diffusion

It is well established that lipid diffusion coefficients in the Lo phase are much lower than those in Ld phase. This is due to the increased lipid order in the Lo phase that results from tighter lipid packing imparted by saturated lipid tail group interactions as well as their interactions with cholesterol.^61^ In our system, the addition of DOPC liposomes causes dissolution of Lo and Ld domains, so there should be an increase in lipid diffusion coefficient after the addition of DOPC liposomes to the SLB patch. To test this, we first made SLB patches by rupturing GUVs with the 2:2:1 DOPC/DPPC/Chol lipid composition, where the Lo and Ld phases were marked with TF-Chol and TR-DHPE, respectively. Then, fluorescence recovery after photobleaching (FRAP) measurements were conducted on ROIs within the Ld domains. After FRAP, DOPC liposomes were added to cause the dissolution of distinct Lo and Ld domains then FRAP was conducted again in the same ROI **(Figure S4)**. As shown in **Figure 8a**, the rate of fluorescence recovery is significantly greater after domain dissolution by addition of DOPC liposomes. Prior to domain dissolution, the diffusion coefficient of TR-DHPE in the Ld phase was 0.50 ± 0.05 µm^2^/s while after dissolution it was 1.83 ± 0.07 µm^2^/s **(Figure 8b)**. The diffusion coefficient measured after domain dissolution is in good agreement with those measured in SLBs composed of DOPC.^61^ Furthermore, the maximum fluorescence recovery, which represents the fraction of lipids that are diffusionally mobile, increases after the dissolution of the domains. These results suggest that there is a wholesale reorganization of the membrane after DOPC liposome addition, rather than just a scrambling of fluorescent probe localization, underscoring the ability of our approach to modulate SLB patch composition.

**Figure 8:**
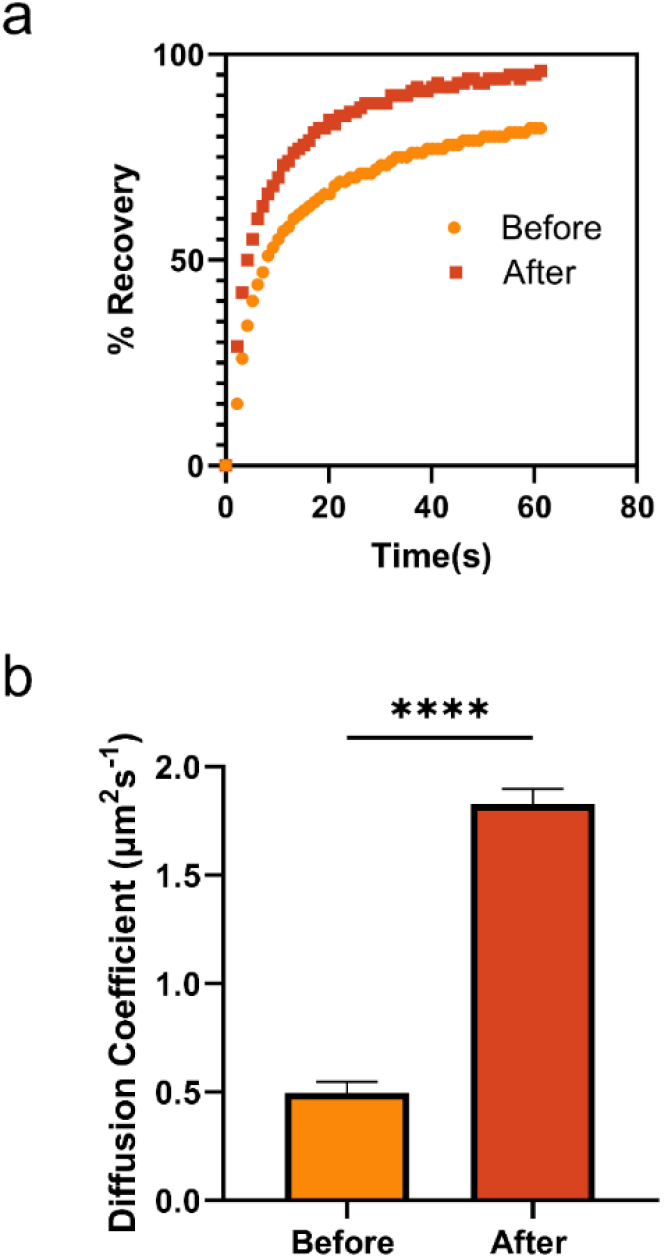
FRAP analysis before and after domain dissolution by DOPC liposomes. (a) Normalized fluorescence recovery curves. (b) Mean lipid diffusion coefficients before and after domain dissolution by DOPC addition. The error bars represent standard deviations, \*\*\*\**p*<0.001.

### Matched SLB Patch and Liposome Compositions

Since addition of DOPC liposomes to phase-separated SLB patches with the 2:2:1 DOPC/DPPC/Chol composition caused Lo and Ld domains to disappear via shifts in overall lipid composition, we predicted that if we added liposomes with the same composition as the SLB patch, the distinct Lo and Ld domains would persist, even as the SLB patch area grew. In this experiment, we started with SLB patches prepared by bursting GUVs with the 2:2:1 DOPC/DPPC/Chol (composition *iii* in **Figure 1**), then we added 100 µL of 0.1 mg/mL of liposomes with the same 2:2:1 DOPC/DPPC/Chol composition and observed changes to the SLB patch, which are shown in **Figure 9** and **Movie S4**. At early time points (t = 100 s) an obvious spreading of TF-Chol fluorescence is observed in accordance with the overall SLB patch growth, along with a slight enlargement of the Ld domain marked with TR-DHPE. As time progresses, the TF-Chol fluorescence spreads over the entire image frame and the Ld domain continues to enlarge. By the end of image collection at 1200 s, the distinct Ld domain is still present, although it has enlarged significantly since the addition of the liposomes. The TF-Chol fluorescence intensity still remains at its lowest level inside the Ld domain, but the contrast between this area has diminished. This indicates that the concentration gradient of TF-Chol is not as sharp as it initially was. However, it is clear that by adding liposomes with a composition that matches that of the SLB patch, the patch can be enlarged without entirely disrupting long-range lipid order that was initially present.

**Figure 9:**
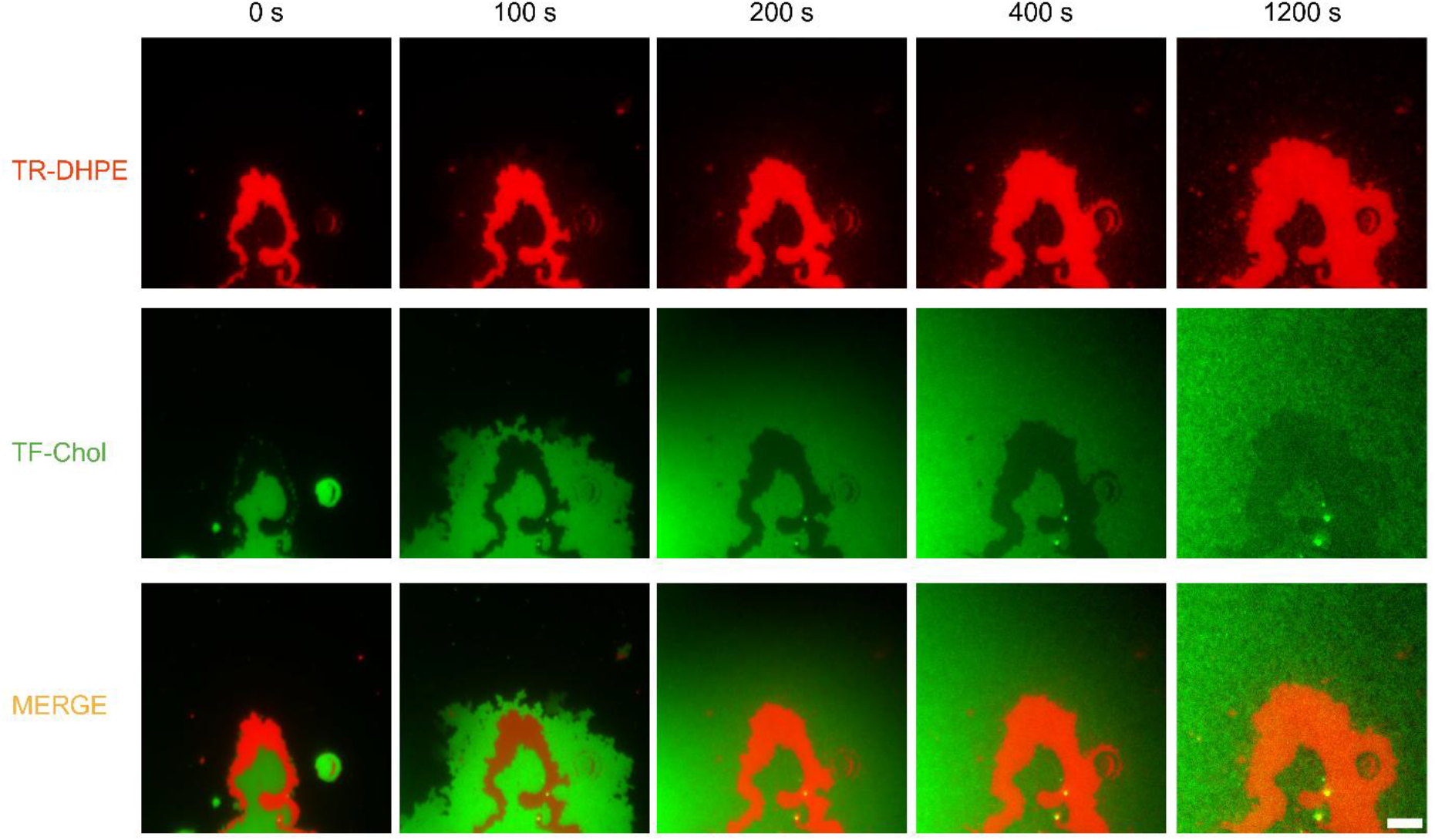
2:2:1 DOPC/DPPC/Chol liposome interaction with 2:2:1 DOPC/DPPC/Chol phase-separated SLB patch. Upon addition of the 2:2:1 DOPC/DPPC/Chol liposomes, the bilayer grows, with both the Lo and Ld phases increase in size, while the membrane two-phase coexistence is maintained. The top, middle, and bottom images represent the Ld (red) labeled with TR-DHPE, Lo (green) labeled with TF-Chol, and the merger of the two channels. The corresponding movie is in **Movie S4** (Supporting Information). Scale bar: 5 µm.

### SLB Patch and Liposomes with Equivalent Cholesterol Fractions

So far, we have shown that addition of DOPC liposomes caused domain dissolution in 2:2:1 DOPC/DPPC/Chol SLB patches, while addition of 2:2:1 DOPC/DPPC/Chol liposomes to the same SLB patch composition preserves and enlarges distinct Lo and Ld membrane domains. Next, we wondered if holding cholesterol concentration constant between liposomes and SLB patches while varying DOPC and DPPC content would alter membrane domains. In this set of experiments, we formed SLB patches with the 2:2:1 DOPC/DPPC/Chol composition and added liposomes with an 4:1 DOPC/Chol composition. Here, the cholesterol content is 20 mole % for both the SLB patches and the added liposomes. Even though the added liposomes are 80 mole % DOPC, we did not observe domain dissolution upon their addition. **Figure 10** and **Movie S5** show representative results. Cholesterol stabilizes the membrane domain by increasing the packing of the saturated acyl chain, while the “kink” resulting from DOPC unsaturated tail group contributes to the loose packing of the liquid-disordered phase.^62^ The observed enlargement of the two phases suggests that the 20 mol % of cholesterol in the added vesicle is high enough to stabilize the Lo phase packing, while at the same time the 80 mol % DOPC contributes in increasing the net DOPC composition in the Ld.

**Figure 10:**
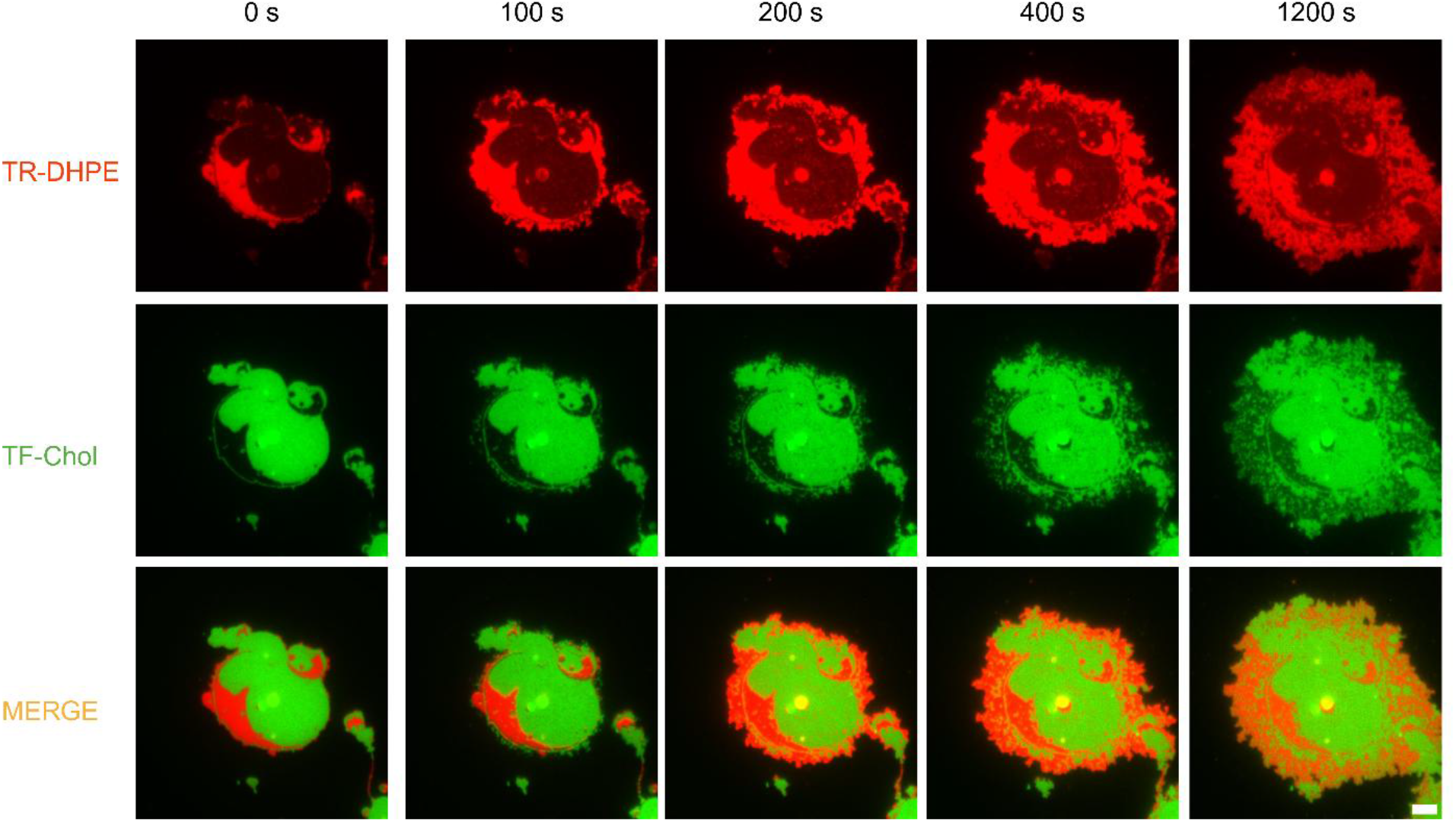
4:1 DOPC/Chol liposome interaction with 2:2:1 DOPC/DPPC/Chol phase-separated SLB patch. The Ld and Lo phases slightly increase in size upon the addition of 0.1 mg/mL 4:1 DOPC/Chol liposomes, while retaining two-phase coexistence. The top, middle, and bottom images represent the Ld (red) labeled with TR-DHPE, Lo (green) labeled with TF-Chol, and the overlap of the two channels. The corresponding movie is **Movie S5** (Supporting Information). Scale bar: 5 μm.

### Redistribution of Lo-binding Proteins

Many membrane proteins are associated with cellular lipid rafts.^63^ Similarly, there are a number of proteins that interact with Lo domain-associated lipids in model membrane systems. A well-known example is cholera toxin B-subunit (CTxB), which binds the ganglioside GM1 with high affinity. Gangliosides, including GM1, are glycosphingolipids that preferentially partition into Lo domains in GUVs and SLBs alike, and fluorescently labeled CTxB is a widely-employed marker of GM1 and thus Lo domains.^64^ Having determined that a fluorescent lipid probe of Lo domains (TF-Chol) becomes homogeneously distributed after domain dissolution upon DOPC liposome addition, we sought to determine if the same was true for a protein bound to Lo domain-associated lipids.

GUVs with a 2:2:1 DOPC/DPPC/Chol with 1 mole % GM1 and 1% TR-DHPE were prepared then allowed to burst on a glass surface to form SLB patches, as previously described. Then the SLB patches were incubated with 10 nM CTxB-Alexa 488 for 30 minutes, before removing unbound CTxB by buffer exchange. At this stage, clear segregation of the CTxB-Alexa 488 and TR-DHPE is observed **(Figure 11, left column)**. If GM1 is omitted from the GUV and SLB patch preparation, CTxB-Alexa 488 does not bind. Next, a 100 µL aliquot of 0.1 mg/mL DOPC liposomes was added to the chamber to initiate the addition of DOPC to the SLB patches. **Figure 11** and **Movie S6** show the time course of TR-DHPE and CTxB-Alexa 488 redistribution. Similar to the case where the Lo and Ld domains are marked with TF-Chol and TR-DHPE, respectively **(Figure 6)**, the addition of DOPC liposomes causes a rise in lipid miscibility, but the rate at which the Lo domains disappear is much slower. For example, at the t = 315 s time point, both the Ld and Lo domains in the vestigial patch are still prominent, even though some of the TR-DHPE and CTxB-Alexa 488 markers have diffused into the nascent SLB patch area. At time points after 720 s, the original patch area is still apparent in the TR-DHPE and CTxB-Alexa488 channels despite the entire ROI being filled with at SLB. This lag in content mixing upon domain dissolution was not observed when the Lo domain was marked with TF-Chol.

**Figure 11:**
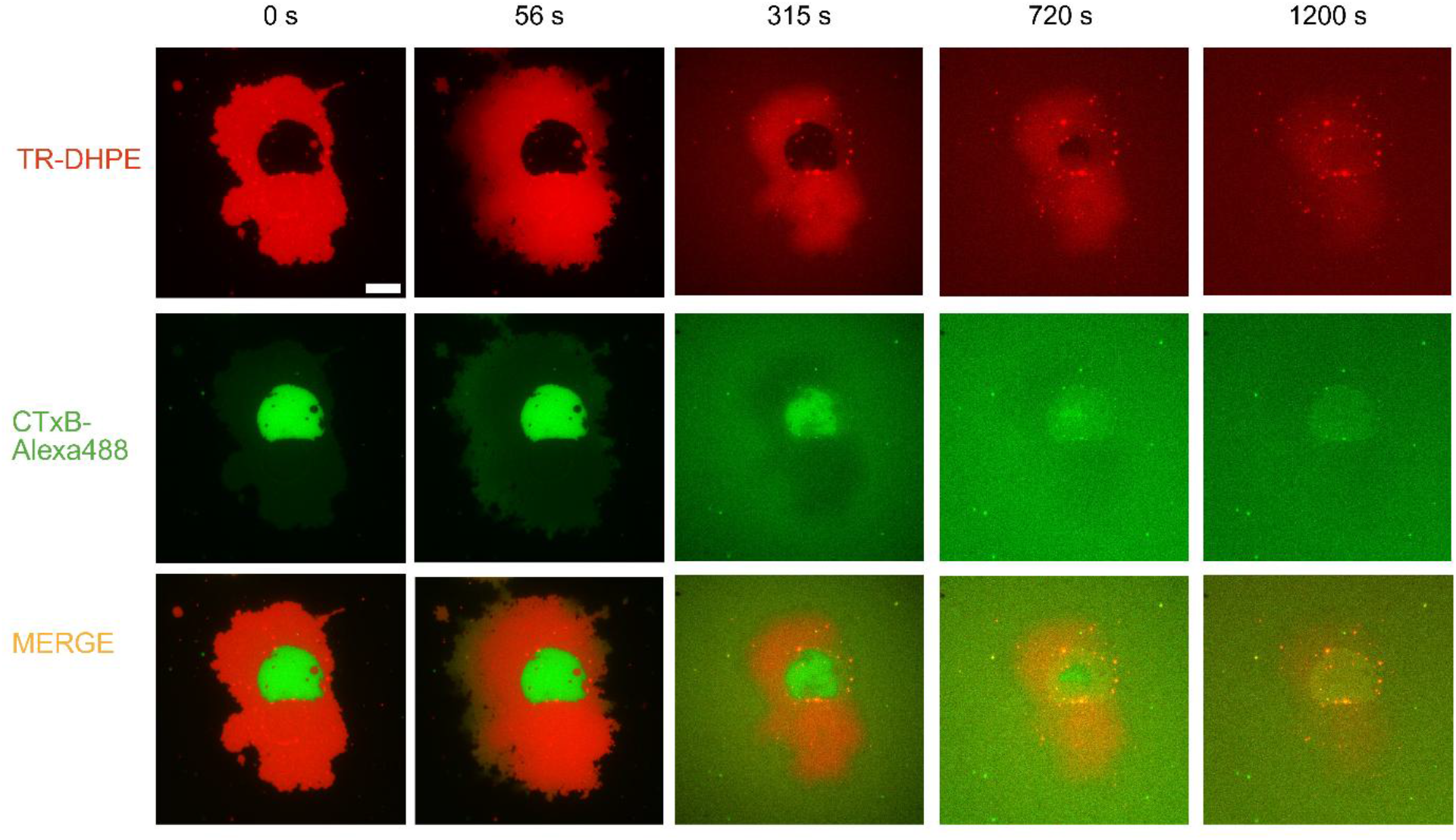
DOPC liposomes induce redistribution of GM1 bound to CTxB in Lo phase. Fluorescence micrographs showing GM1-CTxB Alexa 488 distribution during Lo domain dissolution. The top, middle, and bottom rows represent the Ld (red) labeled with TR-DHPE, the Lo (green) labeled with showing GM1-CTxB Alexa 488, and the merger of the two channels. The corresponding movie is shown in **Movie S6** (Supporting Information). Scale bar: 5 μm

The relatively slow spread of CTxB-Alexa488 fluorescence upon domain dissolution is likely due to a few factors. First, compared to lipid-linked small molecule fluorophores, CTxB has a molecular weight of about 55 kDa, which results in a reduction in its diffusion coefficient relative to phospholipids. In SLBs, typical lipid-linked small molecule fluorophores like TR-DHPE have diffusion coefficients in the range of 1-5 µm^2^/s,^65^ depending on the substrate and background lipid composition. On the other hand, the diffusion coefficient of CTxB bound to GM1 can vary depending on the number of GM1 molecules bound by a single CTxB, which has five GM1 binding sites. The diffusion coefficient of CTxB has been reported to be 0.75, 0.26, or 0.022 µm^2^/s, for CTxB bound to one, two, or three GM1 molecules, respectively.^66^ Clearly, the ability of CTxB to crosslink GM1 molecules that are in close proximity can drastically reduce its apparent diffusion coefficient. Because the GM1 molecules are in closer proximity in the Lo domains in our SLB patches, the crosslinking of GM1s and the resultant reduction in CTxB diffusion coefficient likely explains why the original Lo domain is visible in the CTxB channel after > 20 minutes of DOPC addition to the patch.

## CONCLUSION

Membrane microdomains and lipid rafts have attracted many studies aimed at better understanding their biophysical properties. The dynamic composition of these domains compared to their surrounding membrane makes them rich sites for several cellular processes, including lipid trafficking, signal transduction, protein sorting, and endocytosis. Various studies have also confirmed their association with several physiological conditions such as cancer, neurodegenerative diseases, infectious diseases, and cardiovascular diseases.

The dynamics of membrane microdomains are commonly studied in model systems, such as phase-separated GUVs with predefined lipid compositions. In this work, we used SLB patches with ternary mixtures of DOPC, DPPC, and Chol, with mol percentages of 19:55 25, 2:2:1, and 54:30:16, corresponding to varying area fractions of coexisting liquid-ordered (Lo) and liquid-disordered phases labeled with Chol-TF and TR-DHPE, respectively. We showed that DOPC liposomes added to the SLB patch edge can disrupt and dissolve distinct membrane domains, resulting in a homogeneous phospholipid bilayer. Conversely, a control experiment with liposomes containing 2:2:1 DOPC/DPPC/Chol and 4:1 DOPC/Chol interacting with a 2:2:1 DOPC/DPPC/Chol phase-separated GUV instead shows progressive enlargement of both the Lo and Lo phases while maintaining membrane heterogeneity. We also observed that the domain dissolution induces the redistribution and migration of Lo phase-localized molecule GM1 even after binding with CTxB.

Our approach could offer a route to sequentially tune the composition of SLB patches through the serial exposure of SLB patches to liposomes of varied composition. Doing this would allow the stepwise exploration of multiple regions in compositional space. This could be advantageous for examining lipid composition-dependent protein binding or for interrogating membrane-materials interactions.

## SUPPORTING INFORMATION

Fluorescence images of SLB patch growth by liposome addition, additional fluorescence images of domain dissolution in phase-separated SLB patches upon addition of DOPC from liposomes, fluorescence recovery after photobleaching images obtained before and after DOPC addition to phase-separated SLB patches (PDF).

Movies corresponding to the still images in Figure 6 (Movie S1), Figure S2 (Movie S2), Figure S3 (Movie S3), Figure 9 (Movie S4), Figure 10 (Movie S5), and Figure 11 (Movie S6) (MP4).

## AUTHOR INFORMATION

## Author

Adeyemi T. Odudimu − Department of Chemistry,

Lehigh University, Bethlehem, Pennsylvania 18015,

United States.

## Note

**All authors declare no competing financial interest**

## ACKNOWLEDGEMENTS

This work was supported by a grant from the National Science Foundation awarded to N.J.W (Award number 2044792)

