## Supporting Information for "Domain Dissolution in Supported Lipid Bilayers Triggered by Unsaturated Phospholipid Addition"

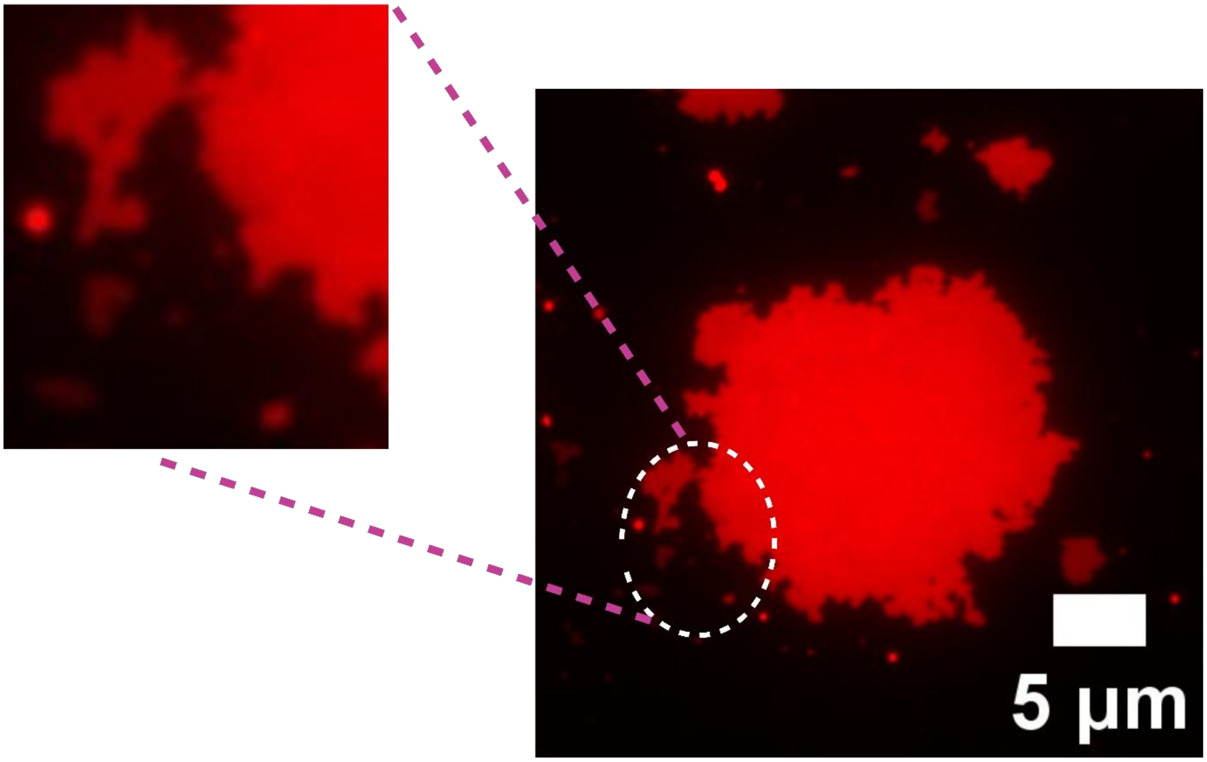


**FIGURE S1**: Formation of fractal-like SLB growth on addition of unlabeled DOPC liposome to a single-phase SLB patch of DOPC/TR-DHPE 99.5:0.5 mol %. Scale bar: 5 µm.

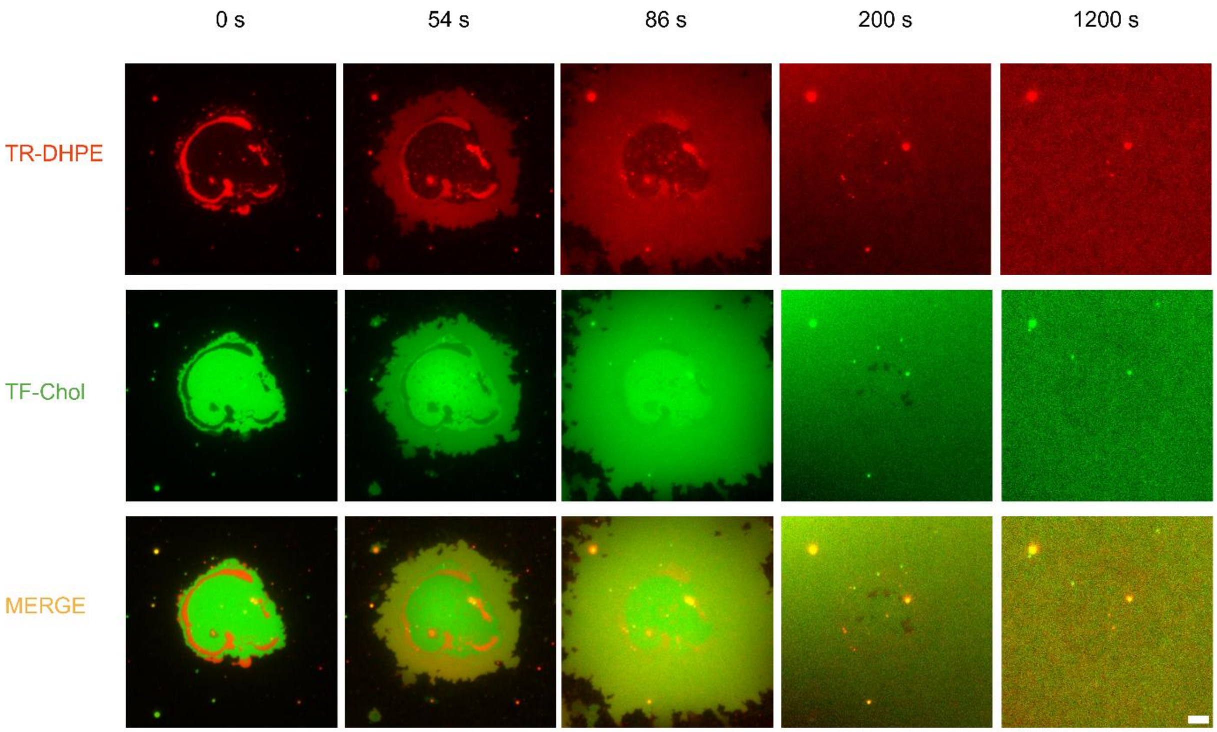


**Figure S2:** Time-lapse fluorescence imaging of 100% DOPC liposome interaction with 19:55:25 mol% DOPC/DPPC/Chol phase-separated GUV, corresponding to composition *iv* in **Figure 2** of the main text. Change in the morphology of the co-existing Lo-Ld phase of phase-separated GUV of 19:55:25 mol% DOPC/DPPC/Chol to form a single phase supported bilayer when 0.1 mg/ml 100% DOPC liposome. The top, middle, and bottom images represent the Ld (red) labeled with TR-DHPE, Lo (green) labeled with TF-Chol, and the overlap of the two channels. The corresponding movie is **Movie S5** (Supporting Information). Scale bar: 5 μm.

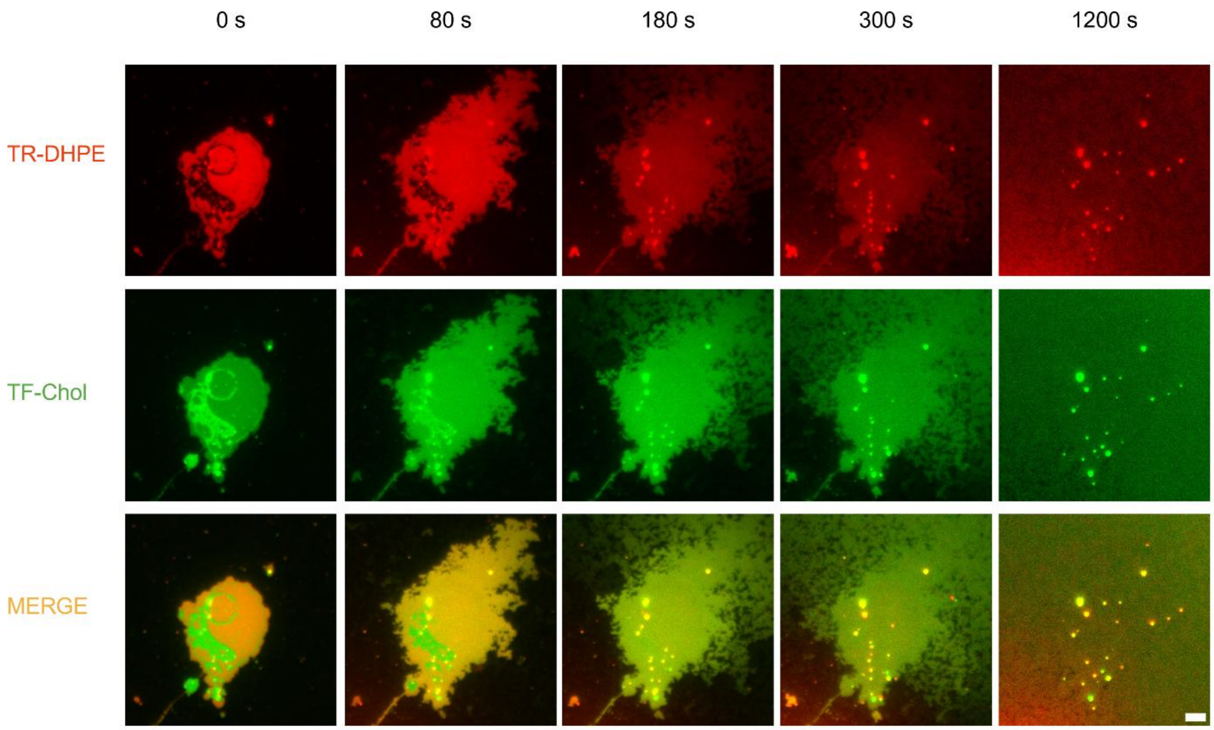


**Figure S3:** Time-lapse fluorescence snapshot of 100 % DOPC liposome interaction with a 54:30:16 mol% DOPC/DPPC/Chol phase-separated GUV, corresponding to composition *ii* in **Figure 2** in the main text. Change in the morphology of the co-existing Lo-Ld phase of phase-separated GUV of 54:30:16 mol% DOPC/DPPC/Chol to form a single phase supported bilayer when 0.1 mg/ml DOPC liposome was added. The top, middle, and bottom images represent the Ld (red) labeled with TR-DHPE, Lo (green) labeled with TF-Chol, and the overlap of the two channels. The corresponding movie is in **Movie S6** (Supporting Information). Scale bar: 5 µm.

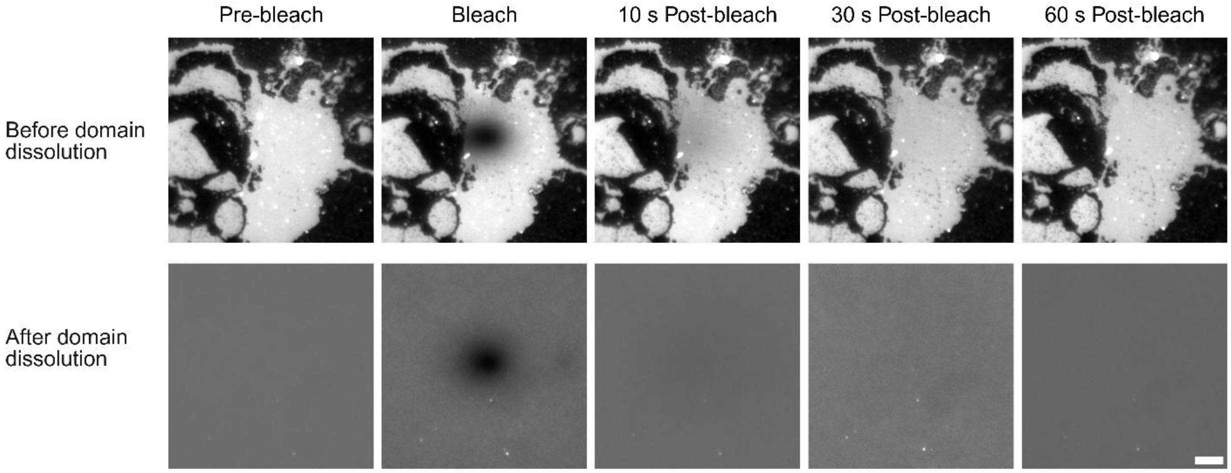


**FIGURE S4:** Fluorescence Micrographs of FRAP analysis of the Ld phase before and after domain dissolution with DOPC liposome. The corresponding diffusion coefficient and % recovery are shown in **Figure 8** of the main text.

**Supporting Movie Captions**

**Movie S1:** Epi-fluorescence microscope movie showing the time-lapse disappearance of membrane microdomain on the addition of 100% DOPC liposomes to a phase-separated GUV-supported bilayer of 2:2:1 DOPC, DPPC, and Chol. There are three movies in parallel. On the left is the TRITC channel (in red) with the Ld phase labeled with TR-DHPE; in the middle is the FITC channel (in green) showing the Lo phase marked with TF-Chol; and on the right is the overlay of the red and green channels. 0.1 mg/mL of 100% DOPC liposomes were added at 0 sec, and the change in the SLB was monitored for 20 minutes, with a 500 msec interval and simultaneous switching between the TRITC and FITC channels. The movies were captured at 512 by 512 pixels and processed at 15 frames per second. Scale bar: 5 µm.

**Movie S2:** Epi-fluorescence microscope movie showing the time-lapse disappearance of membrane microdomain on the addition of 100 % DOPC liposomes to a phase-separated GUV-supported bilayer of 19:55:25 mol% DOPC, DPPC, and Cholesterol. There are three movies in parallel. On the left is the TRITC channel (in red) with the Ld phase labeled with TR-DHPE; in the middle is the FITC channel (in green) showing the Lo phase marked with TF-Chol; and on the right is the overlay of the red and green channels. 0.1 mg/mL of the liposomes was added at 0 sec, and the change in the SLB was monitored for 20 minutes, with a 500 msec interval and simultaneous switching between the TRITC and FITC channels. The movies were captured at 512 by 512 pixels and processed at 15 frames per second. Scale bar: 5 µm.

**Movie S3:** Epi-fluorescence microscope movie showing the time-lapse disappearance of membrane microdomain on the addition of 100 % DOPC liposomes to a phase-separated GUV-supported bilayer of 54:30:16 mol% DOPC, DPPC, and Cholesterol. There are three movies in parallel. On the left is the TRITC channel (in red) with the Ld phase labeled with TR-DHPE; in the middle is the FITC channel (in green) showing the Lo phase marked with TF-Chol; and on the right is the overlay of the red and green channels. 0.1 mg/mL of DOPC liposomes were added at 0 sec, and the change in the SLB was monitored for 20 minutes, with a 500 msec interval and simultaneous switching between the TRITC and FITC channels. The movies were captured at 512 by 512 pixels and processed at 15 frames per second. Scale bar: 5 μm.

**Movie S4:** Epi-fluorescence microscope movie showing the time-lapse increase in the area fraction of the Lo and the Ld phase addition of 2:2:1 mole ratio DOPC, DPPC, and Chol liposomes to a phase-separated GUV-supported lipid bilayer of 2:2:1 DOPC, DPPC, and Chol. There are three movies in parallel. On the left is the TRITC channel (in red) with the Ld phase labeled with TR-DHPE; in the middle is the FITC channel (in green) showing the Lo phase marked with TF-Chol; and on the right is the overlay of the red and green channels. 0.1 mg/mL of 2:2:1 DOPC, DPPC, and cholesterol liposomes were added at 0 sec, and the change in the SLB was monitored for 20 minutes, with a 500 msec interval and simultaneous switching between the TRITC and FITC channels. The movies were captured at 512 by 512 pixels and processed at 15 frames per second. Scale bar: 5 μm.

**Movie S5:** Epi-fluorescence microscope movie showing the time-lapse increase in the area fraction of the Lo and the Ld phase addition of 4:1 DOPC/Chol liposomes to a phase-separated GUV-supported bilayer of 2:2:1 DOPC, DPPC, and Cholesterol. There are three movies in parallel. On the left is the TRITC channel (in red) with the Ld phase labeled with TR-DHPE; in the middle is the FITC channel (in green) showing the Lo phase marked with TF-Chol; and on the right is the overlay of the red and green channels. 0.1 mg/mL of the liposomes was added at 0 sec, and the change in the SLB was monitored for 20 minutes, with a 500 msec interval and simultaneous switching between the TRITC and FITC channels. The movies were captured at 512 by 512 pixels and processed at 15 frames per second. Scale bar: 5 μm.

**Movie S6:** Epi-fluorescence microscope movie showing the time-lapse disappearance of membrane microdomain on the addition of 100% DOPC liposomes to a phase-separated GUV-supported bilayer of 2:2:1 DOPC, DPPC, and Cholesterol containing GM1 after CTxB Alexa 488 interaction. There are three movies in parallel. On the left is the TRITC channel (in red) with the Ld phase labeled with TR-DHPE; in the middle is the FITC channel (in green) showing the Lo phase marked with TF-Chol; and on the right is the overlay of the red and green channels. 0.1 mg/mL of 100% DOPC liposomes were added at 0 sec, and the change in the SLB was monitored for 20 minutes, with a 500 msec interval and simultaneous switching between the TRITC and FITC channels. The movies were captured at 512 by 512 pixels and processed at 15 frames per second. Scale bar: 5 μm**.**
